# Identifying Biomarkers of Retinal Pigment Epithelial Cell Stem Cell-derived RPE Cell Heterogeneity and Transplantation Efficacy

**DOI:** 10.1101/2022.11.22.517447

**Authors:** Farhad Farjood, Justine D. Manos, Yue Wang, Anne L. Williams, Cuiping Zhao, Susan Borden, Nazia Alam, Glen Prusky, Sally Temple, Jeffrey H. Stern, Nathan C. Boles

## Abstract

Transplantation of retinal pigment epithelial (RPE) cells holds great promise for patients with retinal degenerative diseases such as age-related macular degeneration. In-depth characterization of RPE cell product identity and critical quality attributes are needed to enhance efficacy and safety of replacement therapy strategies. Here we characterized an adult RPE stem cell-derived (RPESC-RPE) cell product using bulk and single cell RNA sequencing (sc-RNA-seq), assessing functional cell integration *in vitro* into a mature RPE monolayer and *in vivo* efficacy by vision rescue in the Royal College of Surgeons rats. scRNA-seq revealed several distinct subpopulations in the RPESC-RPE product, some with progenitor markers. We identified RPE clusters expressing genes associated with *in vivo* efficacy and increased cell integration capability. Gene expression analysis revealed a lncRNA (TREX) as a predictive marker of *in vivo* efficacy. TREX knockdown decreased cell integration while overexpression increased integration *in vitro* and improved vision rescue in the RCS rats.

## Introduction

The retinal pigment epithelium (RPE) supports the overlying neural retinal photoreceptor cells that initiate vision. RPE cells provide nourishment, phagocytose photoreceptor cell outer segments, recycle components of the visual cycle, absorb scattered light, regulate ionic homeostasis and contribute to the blood-retinal barrier (Sparrow et al., 2010; Strauss, 2005). In age-related macular degeneration (AMD), RPE cells undergo changes in morphology, proteome, and phagocytic capacity, resulting in dysfunction, cell death and vision loss (Gu et al., 2012; Kopitz et al., 2004; Lin et al., 2011; Vives-Bauza et al., 2008). RPE replacement therapies are under development to treat AMD using embryonic stem cell (ESC)- and induced pluripotent stem cell (iPSC)-derived RPE cells, which have shown promise to rescue vision in animal models and in patients in early clinical trials (da Cruz et al., 2018; Mandai et al., 2017; Sharma et al., 2019). Adult RPE stem cell (RPESC)-derived RPE progeny (RPESC-RPE) are another stem cell-derived source of RPE cells for transplantation to treat AMD (Salero et al., 2012). In our previous work, we showed that subretinal transplantation of a suspension of RPESC-RPE cells in the Royal College of Surgeons (RCS) rat can preserve vision in this model of retinal degeneration (Davis et al., 2017). A clinical trial of RPESC-RPE transplantation for non-exudative (dry) AMD is underway (NCT04627428).

An important step to characterize the identity of cell products for transplantation is to describe the extent of cellular heterogeneity. RPE cells have been traditionally considered a homogenous cell population composed of one cell type, but accumulating evidence points towards RPE cellular diversity (Voigt et al., 2019; Xu et al., 2021). Differences in RPE cell behavior, appearance, and gene expression suggest cellular diversity in the native RPE layer and in stem cell-derived RPE preparations(Cuomo et al., 2020; Ortolan et al., 2022; Whitmore et al., 2014). Previously, we showed that transplantation of RPESC-RPE cells cultured for approximately 4 weeks after thaw from the master cell bank (MCB) is more effective at vision rescue compared to earlier and later time points of *in vitro* differentiation. The molecular mechanisms underlying improved effectiveness of transplants at the 4-week stage of differentiation, however, were unknown.

Here we use single cell and bulk RNA sequencing to investigate the heterogeneity of RPESC-RPE cells and the underlying molecular signatures that confer transplantation efficacy. Our bioinformatic analyses revealed a transcriptomic signature that correlated with successful transplantation. Furthermore, the data revealed a remarkable heterogeneity of RPESC-RPE cell identities, each with a distinct gene expression signature, indicating that multiple, previously unrecognized cell subpopulations are present in the RPESC-RPE product, including a subset of cells with potentially improved chance of integrating into the host RPE after transplantation. Finally, we identified a novel long noncoding RNA (lncRNA) TREX as a biomarker of transplant efficacy in the RCS rat. These findings broaden our understanding of RPE cell product identity and critical quality attributes (CQAs) needed to enhance regenerative approaches to treat RPE dysfunction and vision loss.

## Results

### Bulk RNA-seq uncovers a transplant efficacy gene signature

We sought to determine genes associated with efficient RPE cell transplantation by comparing the transcriptome of RPESC-RPE cultures over an eight-week time course, following our earlier work indicating that the 4-week developmental stage of RPESC is more suitable for transplantation than 2-week or 7-8 week cells (Davis et al., 2017). We utilized the same RPE lines from the earlier study with previously established transplantation effects or a transplant status that could be predicted based on the transplant status of neighboring samples in the timeline (lines 228, 229 and 230; Fig. 1A-B). In addition, we isolated RPE from a fourth donor (line 233) for the experiments. During culture of each RPE line, we collected RNA at the 2, 3, 4, 5, 7, and 8-week time points for library preparation and sequencing. The data were then mapped using STAR aligner (Dobin et al., 2013) (Table S1). We accounted for cell line to cell line variance by batch correction using combat-seq (Zhang et al., 2020); (note that the RPE line 233 at the 4-week time point had much lower counts and that timepoint was discarded from subsequent analysis).

**Figure 1.**
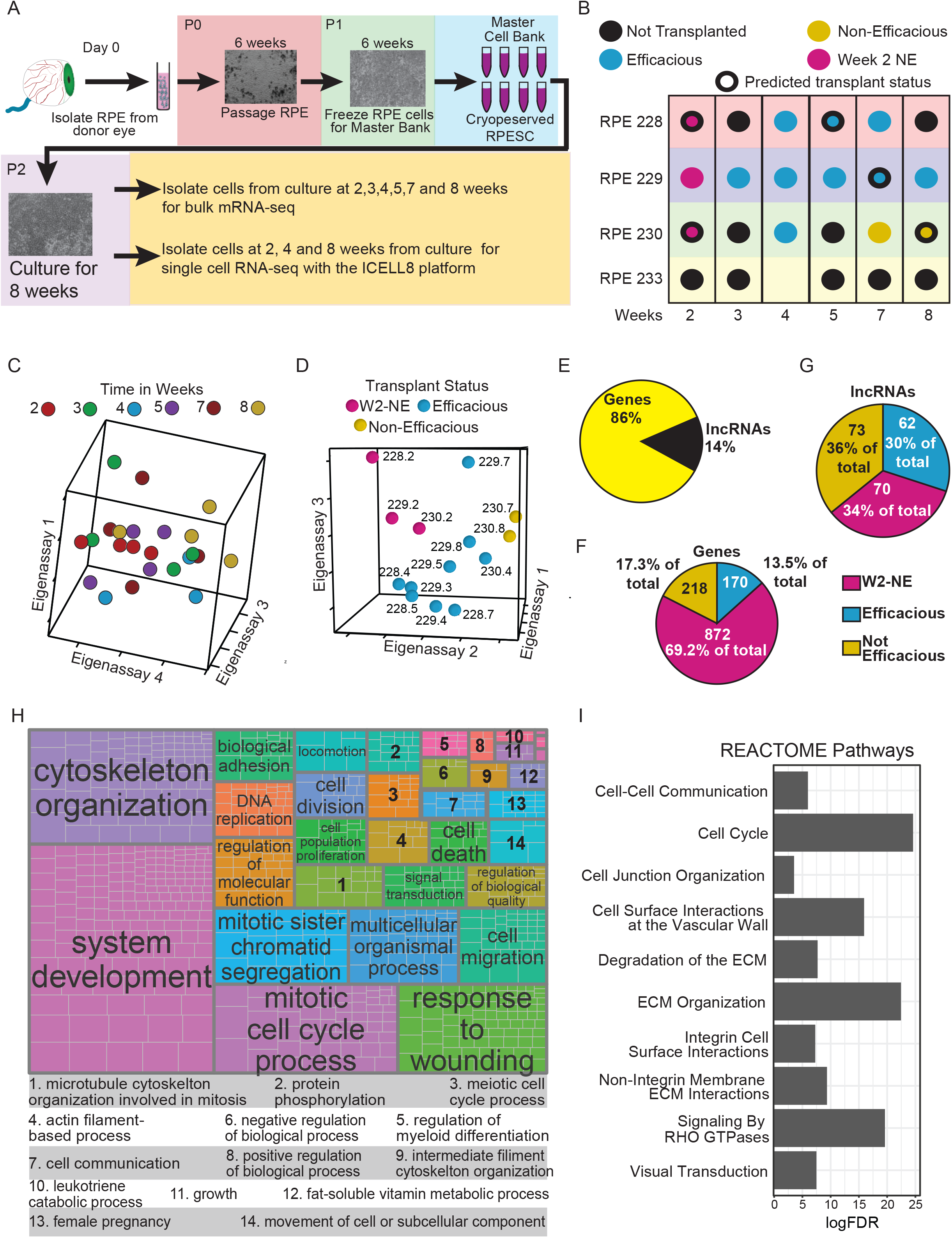
RPESC efficacious for transplant have a distinctive transcriptional profile. (A) Outline of experimental plan for RNA-sequencing. (B) Dot plot illustrating the timepoints RNA was collected, the cell lines used, and the known and predicted transplant status. SVD analysis of the 4000 most variable genes by time (C) and by transplant status (D) was carried out. A discernable pattern was not seen in the time analyzed data, however the data analyzed by transplant status showed a clear separation between groups. (E-G) Distribution of significantly different features by RNA type (E), transplant group for coding gene (F) and long noncoding genes (G). GO enrichment analysis followed by a semantic similarity analysis (Table S3) visualized by treemap (H). Selected pathways from a REACTOME pathway enrichment analysis (Table S4).

We selected the 4000 most variable genes in the whole bulk-RNA-seq dataset and utilized the Singular Value Decomposition (SVD) approach to examine the relationship of samples to each other. Even though previously we utilized weeks in culture as a surrogate measure to separate groups by transplantation efficacy (Davis et al., 2017), the SVD results did not support time in culture as a good variable for grouping samples (Fig. 1C). However, utilizing the known and predicted transplant status of the samples clearly separated them into three groupings based on transplantation status: 2-week non-efficacious cultured RPE cells (W2-NE), Efficacious RPE (EFF-RPE), or Non-Efficacious RPE (NE-RPE) (Fig. 1D). Using this grouping, we proceeded to look for differential gene expression to identify potential biomarkers of transplant efficacy. We used the general linear model approaches in the edgeR and DESeq2 packages to identify differentially expressed genes (DEGs), and included genes identified by both approaches for additional confidence. There were 1465 DEGs with approximately 14% being long-noncoding RNAs (lncRNAs) (Fig. 1E, Table S2). We next determined the extent to which these genes were associated with a transplant group based on their maximum expression and found that the majority of coding DEGs were associated with the W2-NE group (Fig. 1F) while the lncRNA DEGs were associated with each group in roughly equal numbers (Fig. 1G). Proportionately, the lncRNAs made up a much larger fraction of the DEGs in both the EFF-RPE and the NE-RPE. This result is consistent with previous studies on heart cells that highlighted the tissue specificity of lncRNAs; suggesting that these genes may distinguish between groups based on transplant efficacy (Lee et al., 2011).

To gain insight into the biological processes differing between RPE cultures from the W2-NE, EFF-RPE and NE-RPE groups we performed enrichment analysis using the goseq package (Young et al., 2010) (Table S3). Following the GO enrichment testing, semantic similarity analysis (Sayols 2020) was used to group terms (Fig. 1H). Similarly, we performed enrichment for REACTOME pathways (Fig. 1I; Table S4). Some of the most over-represented GO parent terms and REACTOME pathways were associated with proliferation and developmental processes, supporting the concept that the RPESCs are undergoing developmental processes during cell culture and that the cells reach a specific point of intermediate maturation that is ideal for transplantation. However, as demonstrated by our SVD analysis (Fig. 1C,D), time in culture is an imperfect measure of transplant efficacy. One possibility is that as the RPESC cultures develop, multiple subpopulations arise, with one or more subpopulations more effectively conferring transplantation efficacy, and that exactly when those subpopulations arise, and their duration shows variability over time.

### Single cell sequencing reveals changes in RPE heterogeneity over time

We next sought to identify if changes in RPE subpopulations influence transplant efficacy. For these experiments, P2 RPESC cells were cultured for 2, 4, and 8 weeks using the same methodology as for the bulk RNA-seq experiments described above. The ICELL8 platform (Takara) was used to isolate single RPE cells and generate libraries for sequencing with Illumina NovaSeq 6000. Data were processed and mapped with the Cogent NGS Analysis Pipeline (Takara) utilizing the STAR aligner (Dobin et al., 2013). The mapped data was then analyzed using the Seurat (v3) package in R and normalization was performed using the SCTransform pipeline (Hafemeister and Satija, 2019). The dataset was analyzed, and 13 clusters (decreasing in size from 0 to 12) representing subpopulations of RPE cells (Fig. 2A) were discovered. All clusters contained cells from all time points (Fig. 2B), but the subpopulation composition of the whole RPE population changed with time in culture (Fig. 2C). Subpopulations were identified as RPE cells based on expression of previously identified human RPE cell signature genes (Bennis et al., 2015; Strunnikova et al., 2010); all RPE cell clusters demonstrated expression of the majority (136-163/171, 79-95%) of RPE signature genes, underscoring the RPE identity of the subpopulations (Fig. S1).

**Figure 2.**
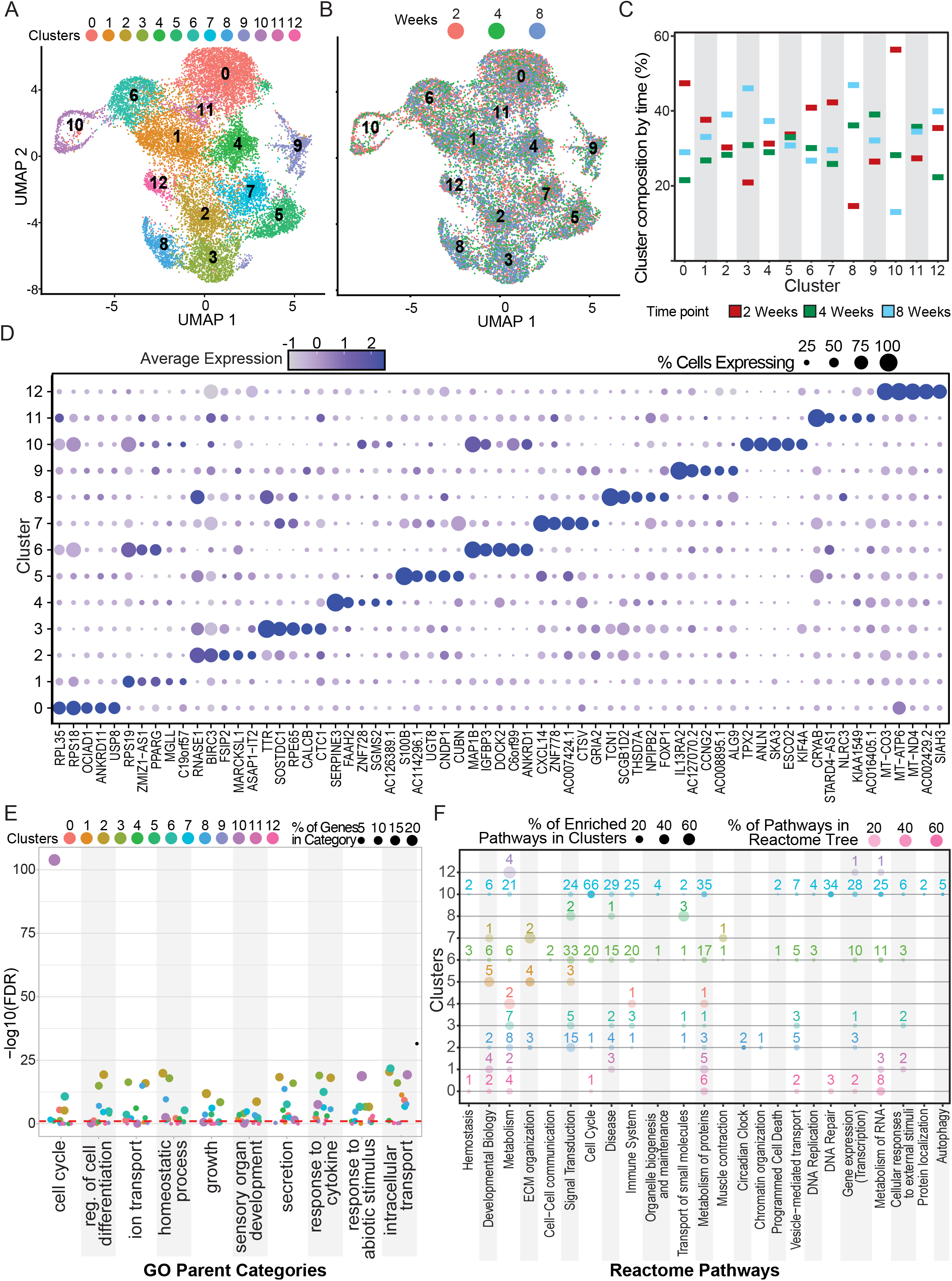
Singe cell RNA-sequencing reveals heterogeneity in RPESC cultures. Single cell RNA-seq (scRNA-seq) was carried out using the well based ICELL8 system with Passage 2 cells collected at 2, 4, and 8 weeks. Data was analyzed using the Seurat package. Dimensionality reduction followed by clustering was used to identify differing clusters of RPESC reveal considerable heterogeneity (A). The clusters appeared to be well mixed when looking at time of collection (B). All clusters had cells from each time period with some clusters showing more variation in their composition (C). Dot plot demonstrating the top five gene markers for each cluster as identified by a Wilcoxon Rank Sum test (D). Using the marker genes for each cluster GO enrichment followed by a semantic similarity analysis (E) and REACTOME pathway analysis (F) was carried out. The numbers above the dots in figure 2F show the number of pathways in each Reactome tree.

To identify genes associated with each cluster, we used the FindAllMarkers function from the Seurat package in R (Fig. 2D, Table S6). Each cluster was associated with between 6 (cluster 11) and 1967 (cluster 10) genes with a median of 300 genes across clusters (Fig. 2D; Table S5). RPE associated genes such as *RPE65* and *BEST1* demonstrated differential expression across the subpopulations. *BEST1*, which has known variable expression levels between central and peripheral retina (Mullins et al., 2007), has higher expression in cluster 2,3,8,9 and *RPE65* showed a similar expression pattern but with increased expression in cluster 7 rather than cluster 9. Transthyretin (*TTR*), which is highly expressed in the RPE of rats and primates (Cavallaro et al., 1990; Pfeffer et al., 2004), also demonstrated variable expression, with cluster 3 having the highest expression followed by cluster 8. *CXCL14*, an immune regulator (Lu et al., 2016) with growth factor activity (Augsten et al., 2009), is highly expressed by RPE cells in the macular region (Whitmore et al., 2014) and demonstrated higher expression in clusters 7 and 5 suggesting a potential macular phenotype for these clusters. Overall, the results support prior work indicating that the RPE layer contains a more heterogenous population of cells than previously considered (Xu et al., 2021).

### Enrichment analysis reveals RPE subpopulation functional specialization

GO and REACTOME pathway enrichment with the hypeR package were utilized to uncover high-level functional differences among the RPE subpopulations (Table S6, S7). The number of enriched GO categories per cluster ranged from 18 categories in cluster 4 to 465 categories in cluster 10, with a median of 76 enriched categories per cluster. After GO enrichments were calculated, a semantic similarity analysis for the enriched terms was performed to group the terms together and assist in visualizing and summarizing the analysis (Fig. 2E; Table S6). For visualizing the signaling pathway enrichments we utilized the Reactome pathways database (v72), which is arranged into multiple hierarchical trees composed of related pathways. For our analysis, we took the top-level terms in the trees and counted the number of enriched pathways under those terms (Fig. 2F; Table S7). The number of enriched pathways for each cluster ranged from 0 in cluster 9 to 329 in cluster 10 with a median of 12 enriched pathways per cluster.

Most clusters showed enrichment for ‘homeostatic processes’, ‘intracellular transport’, and ‘sensory organ development’ GO categories as well as metabolism- and signaling-related REACTOME pathways (Fig. 2E-F; Table S7-S8), as expected given the role of RPE to maintain the retinal microenvironment. Cluster 0 (~20% of cells) demonstrated strong enrichments for metabolic pathways and pathways involved in protein and ion transport. Cluster 1 (~17% of cells) also demonstrated enrichment for pathways associated with metabolism and GO terms associated with metabolism and molecule transport. Cluster 2 (~9% of cells) showed enrichment in a broader number of GO categories ranging from ‘homeostatic process’, and ‘ion transport’ to ‘growth’, ‘neurogenesis’ and a variety of development-related terms. Cluster 2 is enriched for many REACTOME pathways associated with ‘signal transduction’, indicating these RPE cells are likely reactive to environmental challenges and may have a particularly active role in regulating the retinal microenvironment. Cluster 3 (~8% of cells) had a similar enrichment profile to cluster 2, however cluster 3 had less enrichment in developmental pathways and an increased enrichment for metabolic and response to stress functions. In contrast, Cluster 4 (~8% of cells) enriched for fewer GO terms and pathways and the enriched functions were focused on metabolic activities. Together these 5 clusters make up more than 60% of the RPE cells sequenced; they show a spectrum of functions, with cluster 4 being highly metabolic, cluster 2 being highly reactive and clusters 0, 1, and 3 falling between these categories of function.

The pathway enrichments in the remaining clusters also revealed specialization in function. Clusters 5 and 7 showed very similar patterns of enrichment in the GO and pathway analysis for functions associated with locomotion and morphogenesis. The enrichment profile of these clusters could be indicative of subpopulations that could successfully integrate into a RPE monolayer. Notably, cell cycle was substantially enriched in Clusters 6 and 10, indicating these may be proliferative subpopulations. Clusters 6 and 10 were also enriched in signal transduction, and immune and cytokine responses. These clusters are similar to cluster 2 in that they have enrichment profiles with a significant developmental component that could be indicative of populations that benefit cell manufacture and transplantation. Cluster 8 was enriched for functions of molecule transport and stress response. Clusters 9 and 11 were not enriched for any functions and both had few marker genes. Cluster 12, which also had a small number of marker genes, did show enrichment for functions associated with metabolic activities and cation transport. Overall, the RPE subpopulations have overlapping, but distinct functional profiles. Based on the enrichment data, several clusters are candidate subpopulations for efficacious transplantation.

### Intersection of scRNA-seq and bulk RNA-seq data implicates three clusters that are more likely to confer transplantation efficacy

To gain a better understanding of which subpopulations may play a role in transplantation efficacy, we intersected the DEGs correlating positively or negatively with efficiency (Bulk-Eff) from the bulk RNA-seq data with the single cell RNA-seq data. Two sets of DEGs were determined using the FindAllMarkers function in Seurat: the first set was DEGs across the 2-, 4- and 8-week culture timepoints (Time-SC) (Table S8) and the second set were DEGs across the clusters (Cluster-SC) (Table S5). Forty-five percent of the Bulk-Eff DEGs intersected with at least one of the two gene sets from the scRNA-seq data (Fig. 3A). As expected, the Bulk-Eff DEGs were most abundantly expressed in the single-cell data at the 4-week timepoint (Fig. 3B). Importantly, when we compared the Bulk-Eff DEGs to the Cluster-SC data, we found that the genes associated with efficacy were most abundantly expressed in clusters 2, 6 and 10 (Fig. 3C). As noted in the previous section, clusters 2, 6, and 10 were enriched for a variety of functions related to development, environmental reactivity, and, notable in cluster 10, proliferation. This enrichment profile along with the large number of marker genes associated with transplant efficacy make these three subpopulations the lead candidates for playing a key role in successful engraftment and vision rescue after transplantation.

**Figure 3.**
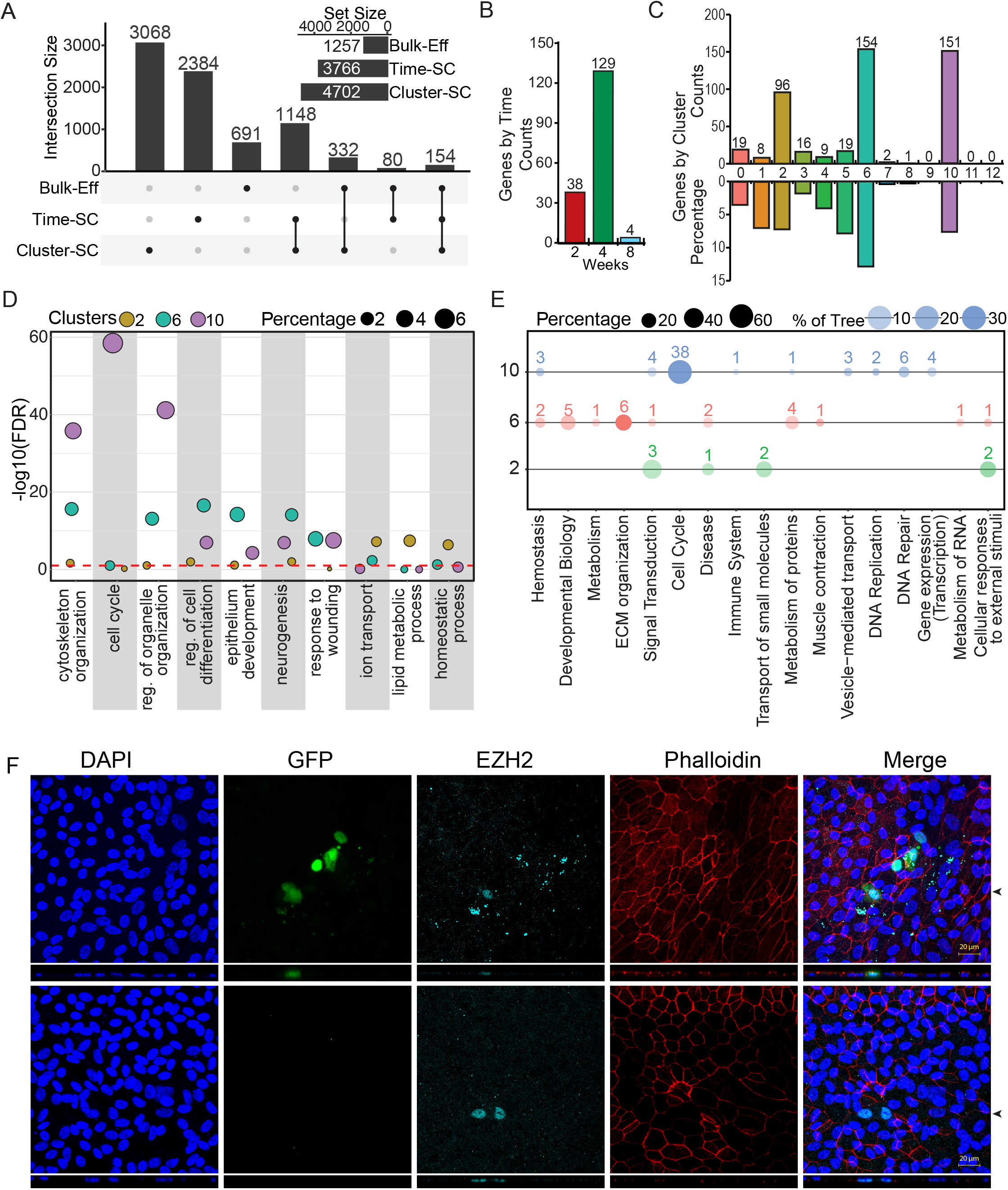
Intersection of Bulk and scRNA-seq data provides insight into potential subpopulations responsible for transplant efficacy. (A) Upset plot showing the overlap between significantly changing genes (DEGs) in the bulk RNA-seq data (Bulk-eff) and the marker genes in the scRNA-seq data based on cluster (Cluster-SC) or time (Time-SC). (B) Membership by time of Bulk-Eff genes and single cell marker genes classified by time in the single cell data. (C) The cluster membership of single cell cluster marker genes found in the Bulk-Eff data by counts and percentage of all marker genes in cluster. We next performed enrichment analysis of the genes shared between the Bulk-Eff and clusters 2,6, and 10. (D) GO enrichment followed by semantic similarity analysis and (E) REACTOME pathway enrichment analysis. (F) RPE cells were immunostained for EZH2 after transplantation of RPE cells cultured for 4 weeks on a RPE monolayer cultured for 8 weeks in an *in vitro* integration assay. EZH2 expression in the monolayer and integrated cells was examined by confocal microscopy. A fraction of integrated cells with GFP expression exhibited EZH2 expression (white arrowheads) indicating that they were members of cluster 10. EZH2 is expressed in the integrating GFP cells (top) and in a small subpopulation of RPE cells in the 8-week-old monolayer (bottom). The boxes below each panel show a section across the z-stack image along the horizontal line indicated by the black arrowhead.

Following up on this discovery, we intersected the markers of clusters 2, 6 and 10 with the Bulk-Eff DEGs and performed GO (Fig. 3D; Table S9) and REACTOME pathway (Fig. 3E; Table S10) enrichment analyses on the intersected gene list for each cluster. The GO analysis revealed some striking differences in the intersected genes from the selected subpopulations (Fig. 3D,E) when compared to the overall cluster GO and REACTOME data (Fig. 2E,F). The intersected gene set from Cluster 10 was highly enriched for terms dealing with proliferation and cell organization. The cluster 2 intersected gene set was enriched for several terms dealing with metabolism and homeostatic processes. On the other hand, the intersected genes from cluster 6 were more specifically enriched for terms related to development and cell differentiation than the total cluster 6 marker genes. In the Reactome analysis, the cluster 2 intersected gene set was enriched for either pathways related to vision or sensing external stimuli. The cluster 6 intersected gene set was enriched for developmental pathways and pathways dealing with the extracellular matrix (ECM), and specifically for pathways interacting with ECM components and for degrading the matrix (Table S9). This cluster 6 enrichment profile may suggest a subpopulation capable of breaking down and integrating into the RPE monolayer. The cluster 10 intersected gene set was again highly enriched for proliferation pathways. Overall, this data analysis reveals that three subpopulations of RPE cells, clusters 2, 6 and 10, have gene expression correlating with properties that could contribute to efficacious transplantation.

### Cluster 10, but not Cluster 2 RPE subpopulations can integrate into an RPE monolayer

The adult RPESC-RPE product is subretinally injected as a cell suspension to enable the possibility of cells integrating into the existing RPE monolayer *in vivo*. As a surrogate of the cell integration process, we developed an *in vitro* assay (Fig. S2). The assay workflow begins by culturing RPE test cells and labeling these with GFP. Then approximately 12,000 GFP+ test cells are plated onto a pre-existing 8-week-old RPE monolayer grown in 24-well Transwell format; at the 8-week stage, the monolayers are highly polarized and exhibit typical mature RPE cobblestone morphology that serves as an *in vitro* model of the native RPE layer. Seven days after plating the GFP+ labeled test cells on the mature RPE monolayer, the cultures are fixed and imaged by confocal microscopy over a pre-set 20 position grid covering approximately 4% of the Transwell surface. GFP+ cells that integrate into the mature GFP-negative monolayer are then counted and the percentage of integrated cells are calculated.

We performed this *in vitro* integration assay by plating 4-week-old GFP labelled RPESC-RPE test cultures on mature monolayers. To determine if integrating cells came from a particular subpopulation, we identified markers for our candidate cluster 2 (YEATS2) and cluster 10 (EZH2) subpopulations and used confocal immunofluorescence to determine if the integrated cells expressed either marker along with GFP. Our results showed that ~90% of EZH2+ cells present in the original suspension had integrated into the RPE monolayer. Cluster 10 only makes up ~3% of the original 4-week RPESC-RPE population, yet EZH2+ cells comprised 22% of all integrated cells (Fig. 3F). This demonstrates that cluster 10 cells will successfully integrate and establish in a preformed RPE monolayer. None of the integrated RPE cells exhibited YEATS2 staining (Fig. S3), indicating that the cluster 2 subpopulation, which represents ~9% of the original isolate, did not successfully establish within the monolayer. These results support our hypothesis that specific RPE subpopulations contribute to an efficacious transplant.

### A long non-coding RNA as a biomarker of efficacious transplantation

After probing for a possible connection between transplantation and changes in the RPE subpopulation composition, we sought to identify a biomarker of efficacious transplantation. Potential efficacy marker candidates were identified by selecting the genes that had a maximum expression of at least 100 counts in EFF-RPE and at least a 2-fold increase in expression over the W2-NE- and NE-RPE groups in our bulk sequencing data. This yielded a total of 36 candidate markers (Table S12). The most consistent and differentially expressed candidate gene is a lncRNA, TCONS_00005049, hereafter referred to as **TREX** (**T**ransplanted **R**PE **Ex**pressed). We examined the level of TREX in the bulk-RNA-seq data of RPESC-RPE samples that had available *in vivo* efficacy data in the RCS rat quantified by optokinetic tracking (OKT) measures of visual acuity. There was a striking positive correlation between OKT data indicating improved vision and increased levels of TREX (Fig. 4A). We next verified TREX levels in these samples by performing qPCR, and again EFF-RPE demonstrated higher levels of TREX (Fig. 4B) than the non-efficacious groups. These data suggest that a threshold level of TREX expression was associated with effective transplantation and improved vision.

**Figure 4.**
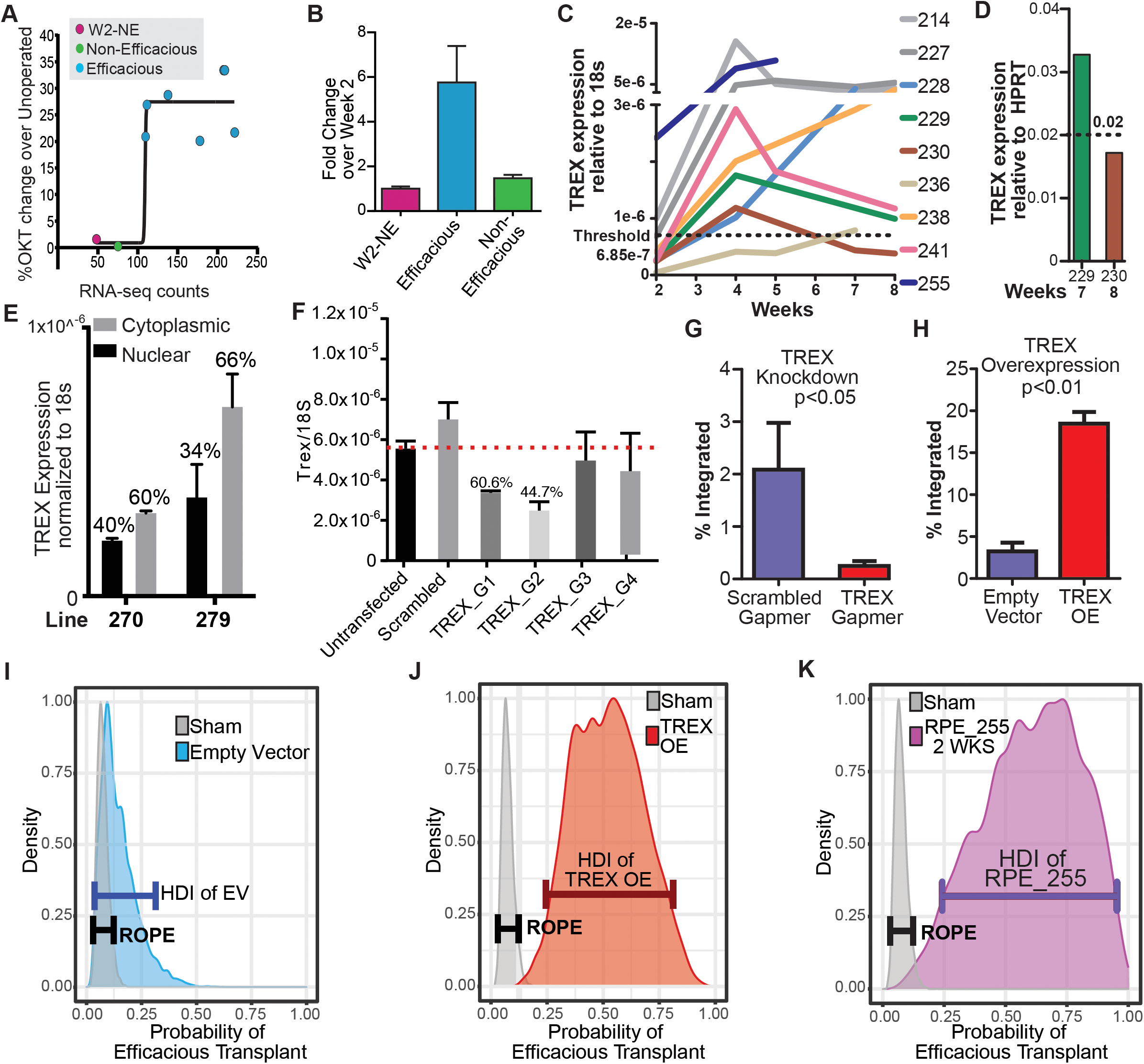
TREX as a biomarker of transplant efficacy. Potential candidates of efficacy were determined by taking the genes that were expressed at least 2-fold higher in the efficacious RPESC cells over the W2-NE and NE RPESC with at least a hundred counts after normalization in the bulk data. (A) A plot of the TREX expression data vs the known OKT results from transplants. (B) Expression of TREX across transplantation groups. (C) TREX expression across multiple lines over time using qPCR with 18s as an internal control. The threshold of TREX expression for a successful transplant is calculated based on the midpoint point the TREX expression of the lowest expressing efficacious cells and highest expressing non-efficacious cells. (D) TREX expression using qPCR with HPRT as the internal control. (E) The cytoplasmic or nuclear fractions of RPE cells were isolated and qPCR was used to look at the distribution of TREX within the cell. (F) Four gapmers against TREX were tested for efficacy in suppressing TREX levels. G2 was used for all knockdown experiments. Results of integration assay (G) for TREX knockdown by gapmer or (H) TREX overexpression (n=3). RPESC transplants were made into the RCS rats and OKT measurements were taken after 90 days. A ‘Region of Practical Equivalence’, or ROPE, was calculated based on the highest density interval (HDI) of 92 sham transplants. The HDI of (I) Empty vector control RPESC, (J) TREX overexpressing RPESC, and (K) RPESC from line 255 having unusually high TREX expression after just 2 weeks were calculated and the densities for each comparison to sham were plotted. The HDI of both the TREX-OE cells and the RPE_255 cells did not overlap with the ROPE demonstrating higher efficacy in transplantation, whereas the empty vector control cells encompassed the ROPE demonstrating no clear difference.

We next undertook qPCR for nine RPE lines including clinical grade cultures obtained under GMP conditions at different cell culture times (Fig. 4C). Most of the lines follow a similar trend with a low level at 2 weeks peaking at 4 weeks. Based on the known transplant data outcomes described previously, we were then able to determine a minimal level of TREX expression (compared to control 18s levels) that was associated with a successful transplant. Note that due to the introduction of proprietary changes made to the 18s taqman probe by its manufacturer, we also established a threshold using HPRT as an internal control for future use, using RPE samples that had demonstrated *in vivo* efficacy (229 at 7 weeks) versus not efficacious (230 at 8 weeks) (Fig. 4D). We next sought to manipulate the level of TREX. As a starting point, we determined if TREX, was primarily in the cytoplasm or the nucleus. Using two different RPE cell lines cultured for 4 weeks, we performed qPCR for TREX on the cytoplasmic fraction or on isolated nuclei and found a 40%:60% distribution respectively (Fig. 4E). Based on this distribution, we decided to use a gapmer-based approach to knock down TREX levels. Gapmer knockdown relies on RNase H to cleave the targets and can be effective in both the nucleus and cytoplasm (Liang et al., 2017). We designed and tested four gapmers against TREX and achieved over 50% knockdown with one of the gapmers compared to scrambled and untransfected controls (Fig. 4F).

To assess the functional effect of TREX levels on RPE transplantation we utilized our *in vitro* integration assay. Using the gapmer based approach, we knocked down TREX in RPESC-RPE cells that had been cultured for 4 weeks and then plated the TREX-knockdown cells on the established mature RPE monolayers used for the integration assay. The scrambled control cells had an integration rate of approximately 2% while significantly fewer TREX-knockdown cells were integrated (~0.2%) (Fig. 4G). We next used a lentiviral overexpression system to overexpress TREX in RPESC-RPE cells cultured for 4 weeks and then performed the integration assay. The empty vector control cells had an integration rate of ~3% whereas the TREX overexpressing cells demonstrated a marked increase in integration with ~18% or 6 times the number of cells integrating into the mature monolayer (Fig. 4H). These results indicate that TREX is not only associated with but is also necessary (Fig. 4G) and sufficient (Fig. 4H) to increase engraftment and transplant efficacy, strongly supporting an important role of TREX to mediate RPE cell integration into a mature RPE monolayer.

Based on our *in vitro* results, we proceeded to assess the role of TREX in RPE transplant efficacy at vision rescue. As previously observed, RPE line 230 cells cultured for 7 weeks were not efficacious after subretinal transplantation in the RCS rat model of retinal degeneration (Fig. 1B). We transfected RPE 230 cells with either an empty control vector or a dox inducible TREX OE virus and cultured them for 7 weeks prior to transplantation. Vision was measured by OKT at 60 days after transplantation (Table S13). Eight rats were transplanted for each condition and five sham (vehicle only) subretinal injections were used as control. The OKT results generally have a bimodal distribution indicative of either a positive effect on vision rescue or a lack of vision rescue. An animal’s vision was considered ‘rescued’ if the OKT measure was above 0.4 cycles/degree and considered ‘not rescued’ if below this threshold. We took a Bayesian approach to the analysis and identified a ‘**R**egion **O**f **P**ractical **E**quivalence’ (ROPE, a range of values representing no effect) based on the highest density interval (HDI, range of values that 95% of the distribution lays under) from the OKT results at p90 of the cumulative sham (vehicle injected) (n=92, Table S13). Using the sham results as a prior probability, we applied a Bernoulli likelihood function to calculate the posterior distribution of the probability of an efficacious transplant of our empty vector control (Fig. 4I) and of TREX OE cells (Fig. 4J). We then sampled from the posterior probabilities to identify the HDI of each condition. In the case of the empty vector control cells, the HDI completely encompassed the ROPE indicating that the control RPE cells performed similarly to the sham as previously observed with the inefficacious 7-week-old RPE 230 cultures. However, in the case of the TREX OE cells, the HDI laid completely to the right of the ROPE, indicating an increase in efficacy associated with RPE cells overexpressing TREX. In addition to our Bayesian approach, we also performed binomial tests for each comparison and found that while the control cells were not significantly different from sham (p=0.09157), the TREX OE cells were different from both sham (p=1.92e-06) and the control cells (p= 0.004227). Hence, there was a positive effect of creating high TREX levels on cell transplant efficacy; we essentially converted a non-efficacious RPE cell preparation to one that effectively rescued vision in this animal model. We next sought to determine if having endogenous high levels of TREX was sufficient to demonstrate transplant efficacy.

RPE line 255 has exceptionally high levels of TREX after only 2 weeks of culture (Fig. 4C), a time point in RPE culture that is usually ineffective at vision rescue compared with 4 weeks of culture prior to transplantation (Davis et al., 2017). We cultured RPE 255 cells for 2 weeks without modification and transplanted them into the RCS rat model at p30, followed by OKT 60 days later. The results demonstrated that the RPE 225 cells with high endogenous TREX were indeed efficacious for transplantation (5 out of 5 animals; Fig. 4K). This result indicates that TREX level was more accurate than time in culture as a biomarker of RPE255 transplantation efficacy. The HDI of the posterior probability of a successful transplant was outside our ROPE and the binomial test also support a significant change (p=1.18e-06). Overall, these results provide strong evidence that TREX is not only a biomarker of efficacious RPE, but that TREX also functions to mediate RPE transplantation efficacy.

## Discussion

The recent expansion in the number and diversity of RPE cell transplant products developed for treating RPE loss in AMD has increased the need to characterize the identity and efficacy attributes of RPE cell products. To advance this knowledge, we performed bulk and single cell sequencing on adult RPESC-derived RPE cell products during *in vitro* differentiation. These studies characterized the individual RPE subpopulations present and identified transplantation efficacy gene markers. Our analysis revealed a clear distinction between the transcriptomes of efficacious and non-efficacious RPE cell products in animal studies that were consistent across different donors, allowing us to identify molecular pathways associated with transplant efficacy and identify potential biomarkers for efficacious RPE cultures.

Our study identified several cell product attributes related to successful transplants by exploring the transcriptional changes in RPESC-RPE cultures between cell populations with varying transplant success in RCS rats. Our earlier work (Davis et al., 2017) showed that time in culture correlated with vision rescue, indicating that time served as a surrogate measure of RPE cell differentiation stage and vision rescue efficacy. Our present transcriptomic analysis demonstrated an incomplete ability to separate the RPE cell transcriptomic profiles based solely on time, pointing out the need to dissect RPE cellular heterogeneity at each timepoint. Recent studies using single cell approaches have reported multiple RPE subpopulations within the stem cell-derived, native human and mouse RPE with unique transcriptional and morphological characteristics, indicating that the RPE cell layer is composed of heterogenous cell populations that may have differing functional and clinical capabilities (Lee et al., 2022; Petrus-Reurer et al., 2022; Voigt et al., 2019; Xu et al., 2021). Identification of the subpopulations responsible for vision rescue can improve regulatory evaluation and improve RPE cell products to result in better transplantation outcomes.

This transcriptomic analysis identified 13 subpopulations within the RPESC-RPE cultures with distinct patterns of gene expression, consistent with recent findings from other laboratories (Petrus-Reurer et al., 2022; Xu et al., 2021). Three clusters, 10, 6 and 2, demonstrated enrichment for pathways associated with cell differentiation and proliferation implying that these subpopulations had progenitor characteristics and were candidates to improve both cell manufacture and transplant success. To enhance our ability to identify subpopulations with more effective transplantation outcomes and uncover potential biomarkers, we developed an *in vitro* integration assay to mimic engraftment after *in vivo* transplantation when transplanted cells insert into the host RPE monolayer. This assay provides a more cost effective and higher throughput method than animal models to examine how an RPE cell population performs in the early engraftment step of a successful transplantation. Additionally, it allows insight into how the genetic manipulation of RPE populations may affect the transplant competency of specific cellular subpopulations. Using the *in vitro* integration assay, we found that a subset of RPE cells were better able to integrate into a mature RPE monolayer. The integrating RPE cells were more likely to express EZH2+, a marker of cluster 10. EZH2 is a polycomb transcription factor involved in histone methylation and stem cell self-renewal and differentiation (Cao et al., 2002; Kamminga et al., 2006; Karantanos et al., 2016). Notably, cluster 10 showed the highest level of cell cycle markers, underscoring its stem- or progenitor-like state. Further study of the prospectively enriched cluster 10 subtype will be worthwhile to understand the roles of this important subpopulation in RPE layer cell biology, RPE cell product manufacture, and RPE transplantation safety and efficacy outcomes. It is possible that interactions between different clusters within the product are beneficial, and we anticipate future studies to alter RPE subtype composition will be needed to determine the optimal product composition.

Our analysis identified potential biomarkers of transplant efficacy, including our top candidate, the lncRNA TREX. Recently, lncRNAs have emerged as potential markers for predicting the quality of other types of tissues and cell products used for transplantation (Wong et al., 2019; Zou et al., 2019). Manipulating TREX expression in our *in vitro* RPE integration assay suggests that TREX may regulate cellular processes involved in engraftment into the host RPE. However, TREX could be utilizing a wide variety of processes ranging from upregulating cell migration to sustaining cell survival to increase RPE potency and further research is needed to understand the mechanism. *In vivo* transplantation experiments support a role for TREX in successful transplantation. An RPE cell line that expressed unusually high levels of TREX was effective at the 2-week stage of differentiation that is usually ineffective. Furthermore, knockdown of TREX caused efficacious REP lines to no longer support cell integration *in vitro*. Finally, over-expression of TREX in non-efficacious RPE lines improved the probability of vision rescue. These results provide strong evidence that TREX is directly involved in mediating transplant efficacy and that TREX can predict the efficacy of RPE cell therapy products. Thus, TREX is a critical attribute and candidate potency biomarker for the RPESC-RPE cell product. While the emphasis of RPE cell product development has been placed on overall RPE purity assessed by canonical RPE markers, it is likely that subpopulation heterogeneity found in adult RPESC-RPE cells is reflected in other RPE cell products such as those derived from pluripotent stem cells (35705015). Our work provides a path to establish a level of TREX expression to identify RPESC-RPE cell products, which may apply to other types of RPE cell products, with the highest likelihood of successful transplantation.

In conclusion, our findings highlight the importance of characterizing at the single cell level stem-cell derived RPE cell products designed for cell transplantation. Knowledge about the identity and quality attributes of RPE cell products is needed to guide their successful development. Our findings highlight the importance knowing the subpopulations present within an overall bulk RPE population. Further in-depth characterization of the different subpopulations of RPE cells present in cell replacement products will be valuable to improve manufacture, regulatory evaluation and transplant efficacy of RPE cell products to benefit retinal patients with degenerative diseases such as dry AMD.

## Methods

### RPE cell culture and subject details

All RPE cell lines were generated from donor eyes obtained from certified eye banks with consent for research use. The donor details are listed in Table S14. RPE cells were cultured in a 24-well plate at 100K cells per well. Different RPE lines were used for different experiments in this study due to the limited availability of adult RPESC-derived RPE cells. RPE cells used in scRNA-Seq experiments were cultured on Transwell inserts (Corning, Corning, NY). Cultures were maintained in RPE medium: DMEM F12 50/50 medium (Corning), MEM alpha modification medium (Sigma-Aldrich), 1.25 mL Glutamax (Gibco), 2.5 mL Sodium Pyruvate (Gibco), 2.5 mL Niacinamide (1M; Spectrum Chemical Inc., CA), 2.5 mL MEM NEAA (Gibco), 2% or 10% heat inactivated fetal bovine serum, supplemented with THT (Taurine, Hydrocortisone, Triiodo-thyronin), and 1.25 mL N1 medium supplement (Sigma-Aldrich). Cells were incubated in a humidified incubator at 37 °C and 5% CO_2_ and the medium was replaced every 3 days.

### Bulk and single cell RNA sequencing preparation

Bulk and single cell RNA sequencing were performed on cultured RPE cells collected at 2-, 4-, and 8-weeks post plating. Single cell suspensions were prepared using 0.25% Trypsin (Thermo Fisher Scientific, Waltham, MA) and processed for either bulk or scRNA sequencing. For bulk sequencing RNA was isolated with Direct-zol RNA kit (Zymo Research, Irvine, CA). Library preparation was then carried out with the TruSeq Stranded Total RNA kit (Illumina, San Diego, CA) ribo-depleted by the University of Rochester Genomics Research Center and sequenced using a NextSeq550 high-output flow cell generating 2 × 151-bp read lengths.

For scRNA-seq, the RPE single cell suspension was stained with 1 μl SYTO64 dye (Invitrogen, Carlsbad, CA) in 1mL PBS for 20 minutes at room temperature, then washed twice in fresh PBS. Next, cells were diluted to 25,000 cells/mL and dispensed into ICELL8 3’ DE chips (Takara Bio, CA) using an MSND device (Takara Bio). Cell dispensing, and in-chip reverse transcription PCR were performed using a 3’ DE Chip and Reagent kit (Takara Bio) according to the manufacturer’s instructions. Following the extraction of PCR products, cDNA samples were concentrated and purified using a DNA Clean & Concentrator-5 kit (Zymo Research) and purified using a 0.6X proportion of AMPure XP magnetic beads (Beckman Coulter, Brea, CA) according to manufacturer protocols. Library preparation was performed using a Nextera XT DNA Library Preparation Kit (Illumina) according to the Takara’s 3’ DE chip and reagent kit instructions. The quantification, and quality checks of the cDNA products were carried out at the University at Albay’s NextGen Sequencing core facility. The concentration of cDNA products was quantified using a Qubit Fluorometer and the Qubit dsDNA HS Assay Kit (Thermo Fisher Scientific). The quality of the cDNA product was checked using an Agilent High Sensitivity DNA Kit and Agilent 2100 Bioanalyzer (Agilent Technologies, Palo Alto, CA) to ensure the complete removal of contaminants. Libraries were sequenced using a NovaSeq 6000 high-output flow cell generating 2 × 150-bp read lengths (GeneWiz).

### Data processing and analysis

For bulk sequencing, the University at Rochester core facility processed the raw Illumina BCL files and provided fastq files. Files were then mapped to hg19 and converted to bam files using STAR (v2.4). Bam files were then read into R using the GenomicAlignments package (code available at https://github.com/neural-stem-cell-institute/RPESC_TREX) to generate a counts matrix for further analysis. For single cell sequencing, raw Illumina read BCL Files were converted to fastq files using the bcl2fastq2 software (bcl2Fastq v2.19.1, Illumina, Inc). For scRNA-seq the, fastq files were then merged into read1 (ICELL8 barcode sequence) and read2 (transcript sequence) fastq files. Using the metadata file from the ICELL8 system containing single cell well information their specific nanowell barcodes, read1 and read2 files were demultiplexed based on nanowell barcodes. The sequence reads were then converted to bam files and mapped to hg19 using STAR (v2.5). The code utilized for processing the ICELL8 data can be found at github (https://github.com/neural-stem-cell-institute/sc-pipeline). The final transcript read counts RData file was used as the input reads matrix for further analysis in R.

All code as well as package versions for the analysis can be found at https://github.com/neural-stem-cell-institute/RPESC_TREX. Briefly, bulk data was analyzed with the EdgeR and DESeq2 packages to identify differentially expressed genes. Single cell data was analyzed with the Seurat package (V3.1). Strict criteria were used for the QC of the single cell dataset, including a minimum feature cutoff of 200 and a minimum cell cutoff of 3. Different clusters were identified according to Seurat’s clustering workflow. scRNA-seq enrichment analysis was performed with the screp package (https://github.com/neural-stem-cell-institute/screp/)

### Lentiviral Vectors Production and Infection

The exonic TREX sequence (TCONS_00005049) was downloaded from the UCSC genome browser (hg19) and a TREX insert with 15bp overhangs was synthesized with GeneArt Gene Synthesis (Thermo Fisher Scientific). The insert was then cloned into a EcoRI cut TetO-FUW vector (Addgene) using an In-Fusion HD Cloning kit (Takara Bio) and sequenced (GeneWiz) to verify plasmid. Lentivirus was generated in 75% confluent 293FT cells by co-transfecting the packaging plasmids pCMV-pLNV and pCMV-pVSVG, as well as either the TetO-FUW-TREX or TetO-FUW plasmids into the cells with the XtremeGene HP DNA transfection reagent at a ratio of 1:2.5 according to manufacturer’s protocol. Supernatant was collected 24 and 72 hours after transfection followed by centrifugation at 21,700g for 2.5 hours to concentrate the viral particles. Viral particles were tittered via qPCR (ABM qPCR Lentivirus Titration Kit) before storage at −80°C.

### Animal maintenance, transplantation, and analysis

Royal College of Surgeons (RCS) rats were obtained from Dr. Shaomei Wang and Long Evans rats from Taconic Biosciences, Inc. were maintained under a 12-hour light/dark cycle according to IACUC-approved procedures. Transplantations were carried out as previously described (Zhao et al., 2017). Briefly, RCS rats at P28–P32 days were treated with cyclosporine (210 mg/L), then a 33-gauge needle was used to inject 1.5 μL of RPESC-RPE cell suspension or BSS vehicle control under the retina under isoflurane anesthesia. Surgical success was confirmed by visualization of a subretinal bleb using optical coherence tomography.

The spatial frequency threshold for opto-kinetic tracking (OKT) was measured (Table S1s) by observers masked to treatment group using a device (CerebralMechanics) and methods previously described (Douglas et al., 2005; Prusky et al., 2004) (Table S13). The results were characterized by a non-normal bimodal distribution leading us to assign transplants as either efficacious or not efficacious. In addition to binomial testing, a Bayesian approach was utilized to compare groups using R. The code for the analysis is available at https://github.com/neural-stem-cell-institute/RPESC_TREX. Briefly, a region of practical equivalence (ROPE) was established using the results from 92 sham experiments. Each group was then compared to this ROPE to determine if a treatment demonstrated an unambiguous difference to the sham experiments.

### In vitro Integration assay

A ‘receiver’ monolayer of RPE (RPE line 270) was grown on Transwells inserts (6.5 mm diameter) for 8 weeks. ‘Donor’ RPE cells at P2 (transfected with a GFP lentiviral maker) were grown for 4 weeks, made into a single cell suspension, and 12,000 RPE cells were transplanted on to the ‘receiver’ RPE monolayer. After one week, Transwells were washed, fixed with 4% paraformaldehyde, and imaged using an LSM780 Zeiss confocal microscope (Zeiss, Germany). Twenty images, whose locations were distributed across the Transwell and in the same relative position to each other, were taken for each Transwell. The images covered ~4% of the wells surface. Confocal images were analyzed using the Zeiss ZEN software (V3.1). Integrated cells were identified by determining if a nucleus from a ‘donor’ cell was in the same z-plane as ‘receiver’ nuclei using the 2.5D view. The percent of integrated cells for each sample was calculated using the following equation:

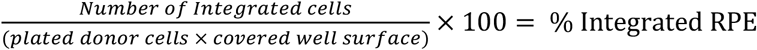

### Immunofluorescence staining

Fixed Transwell membranes were immunostained for EZH2 and YEATS2 to determine the identity of integrated cells. Cells were permeabilized using 1% Triton X-100 in DPBS for one hour at room temperature. After removing Triton X-100 and rinsing 2 times with DPBS, cells were incubated with EZH2 and YEATS2 primary antibodies (Thermo Fisher Scientific) according to the manufacturer’s instructions. Next, primary antibodies were removed and cells were washed with DPBS twice and incubated with secondary donkey anti rabbit or goat anti mouse antibody conjugated with Alexaflour 647 or Alexaflour 555 dyes and DAPI (1:1000) diluted in DPBS with 0.5%bovine serum albumin (BSA) for 1 hours at room temperature. Immunostained cells were then washed 2 times with DPBS for 5 minutes at room temperature and mounted on glass slides using Fluoromount G (Thermo Fisher Scientific) and a coverslip. Samples were imaged using an LSM780 Zeiss confocal microscope (Zeiss, Germany).

### Quantitative RT-PCR

RNA was isolated with the Direct-zol RNA kit (Zymo Research) and cDNA was generated using the Superscript VILO kit (Thermo Fisher Scientific) according to manufacturer’s instructions. Quantitative RT-PCR was performed using TaqMan gene expression assays and TaqMan Universal Master Mix (Thermo Fisher Scientific). Either 18s-VIC or HPRT-VIC was used as an internal control for all reactions.

## Acknowledgements

This study was funded by the National Eye Institute, National Institutes of Health (R01EY029281 to J.H.S.). We thank Malcolm Moos, Ph.D. for useful discussions about single cell strategies to characterize RPE cells. We are grateful for the donors and the Tampa Lions Eye Institute for Transplant & Research, Tampa, Florida, the Eye Bank for Sight Restoration in NY, NY, and National Disease Research Interchange for providing the tissues that were essential for this research.

## Author Contributions

Conceptualization, S.T., J.S. and N.C.B.; Methodology, F.F., J.M.,S.T., J.S. and N.C.B.; Software, F.F. and N.C.B. Validation, F.F.,J.M. and N.C.B. Formal Analysis, F.F., Y.W.,A.L.W.,S.B. and N.C.B.; Transplantation and Analysis, J.M., C.Z, N.A., G.P., N.C.B.; Investigation, F.F., J.M.,Y.W.,A.L.W.,C.Z.,S.B. and N.C.B.; Resources, S.T., J.S. and N.C.B.; Writing – Original Draft, F.F. and N.C.B.; Writing – Review & Editing, F.F., S.T., J.S. and N.C.B.; Visualization, F.F., Y.W., A.L.W. and N.C.B.; Supervision, S.T., J.S. and N.C.B.; Project Administration, J.S. and N.C.B.; Funding Acquisition, J.S. and N.C.B.

## Materials and Correspondence

Further information and requests for resources and reagents should be directed to and will be fulfilled by the Lead Contact, Nathan Boles.

## Data Availability

The raw scRNA-Seq data generated during this study are available at GEO; GSE211189 (https://www.ncbi.nlm.nih.gov/geo/query/acc.cgi?acc=GSE211189).

## Code Availability

Code written for this study is available at github (https://github.com/neural-stem-cell-institute/RPESC_TREX).

---

**Figure S1.**
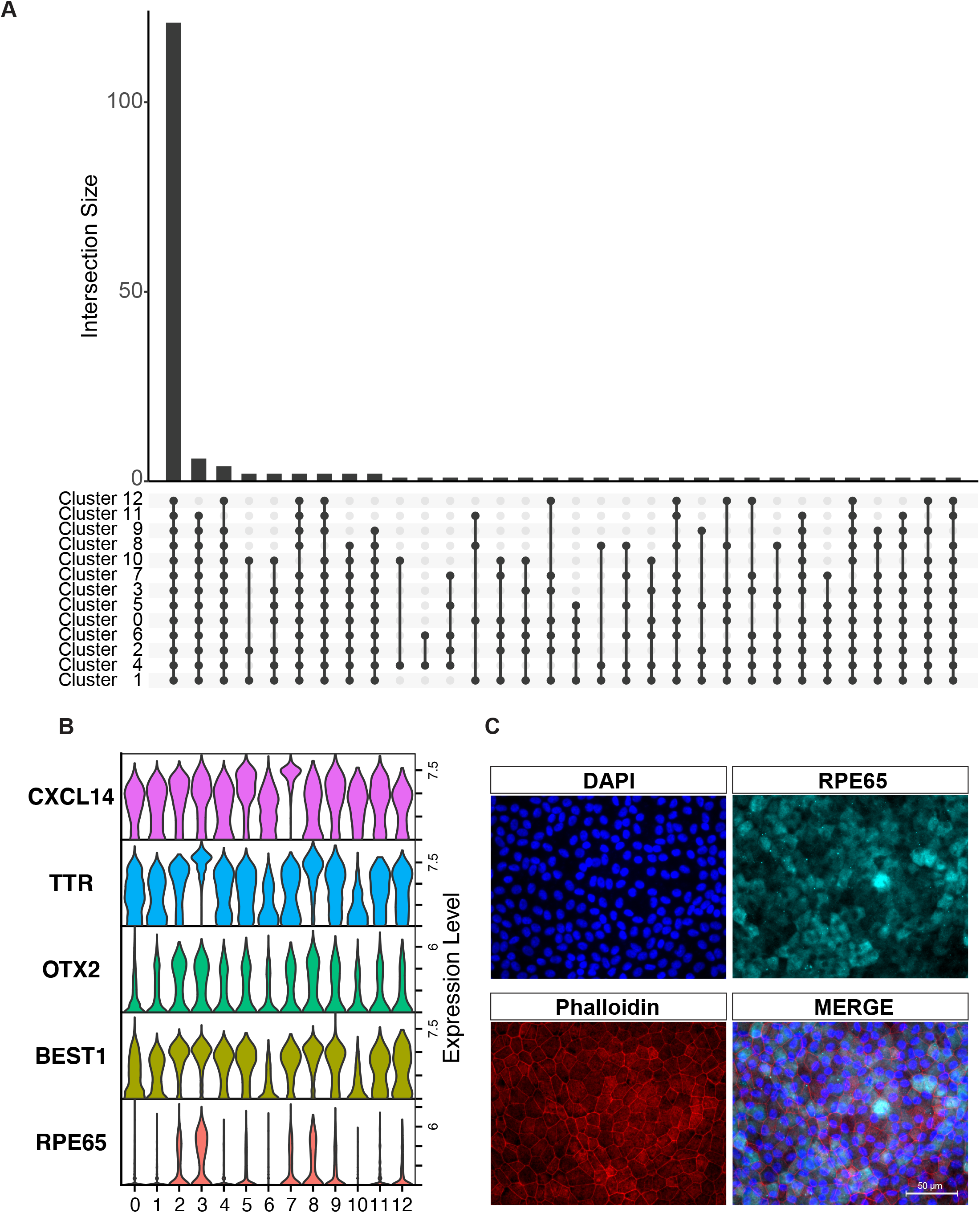
Expression of RPE related genes. (A) Intersection of RPE signature genes with genes expressed in each cluster. (B) Select gene expression using simplified violin plots. (C) Expression of RPE65 in 8-week-old RPE cultures.

**Figure S2.**
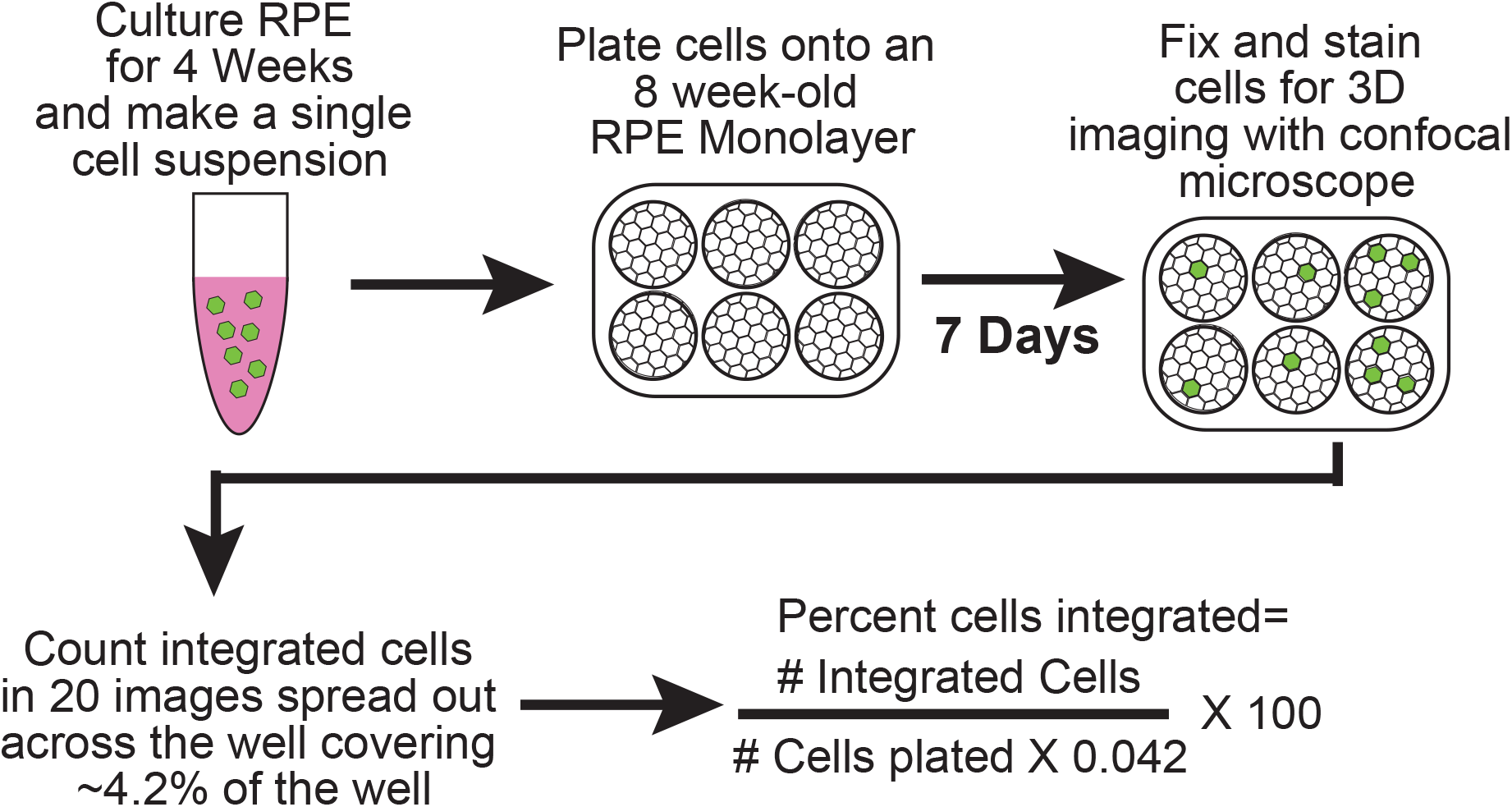
Outline of Integration Assay. Description of novel integration assay’s experimental design and analysis method.

**Figure S3.**
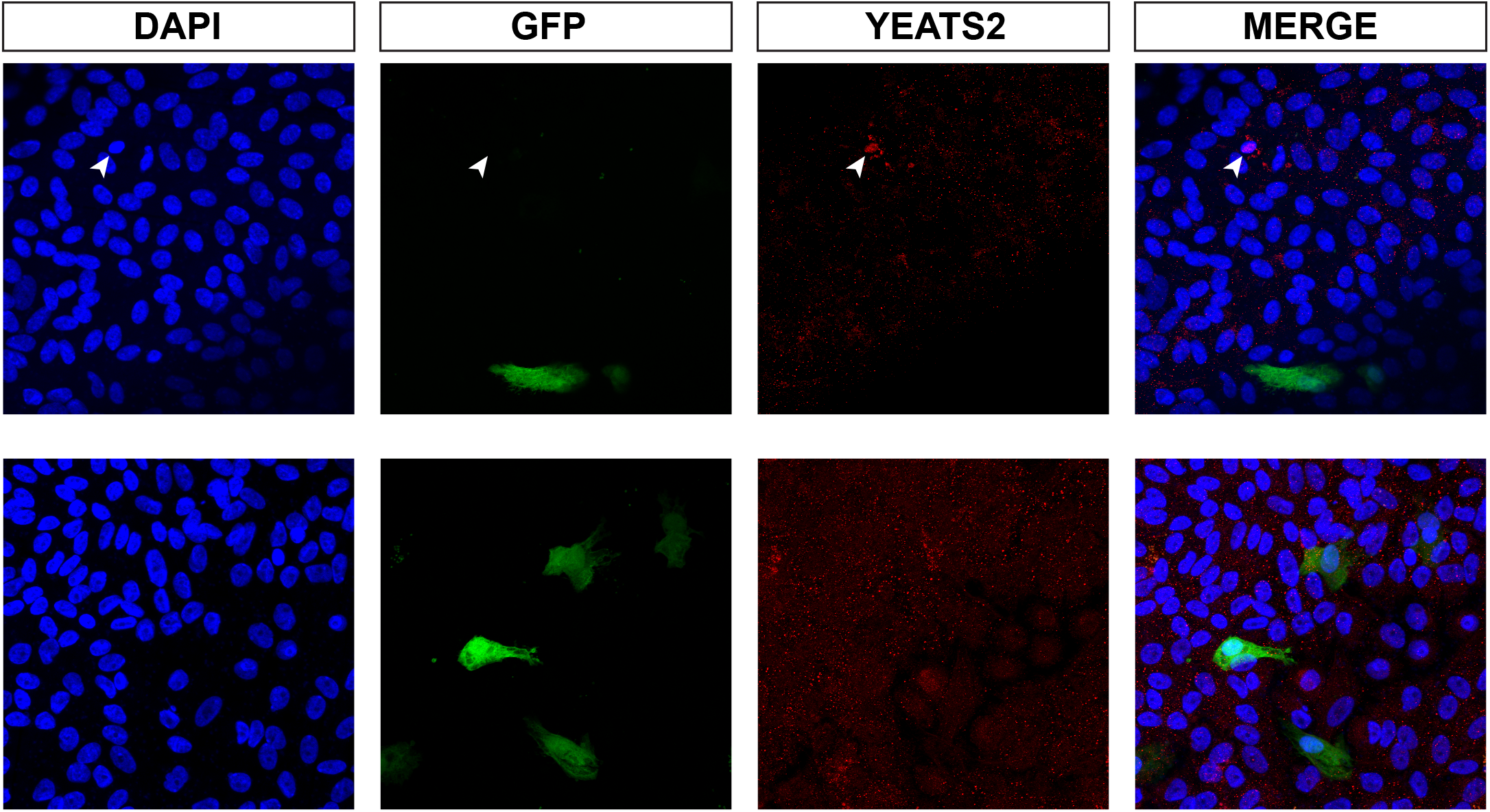
Expression of YEATS2 in RPE cells. YEATS2 was expressed in a subpopulation of 8-week-old RPE cells cultured on Transwells (top row, white arrowheads), but no YEATS2 expression was found in integrated RPE cells labeled with GFP (bottom row).

